# RyR2-Dependent Calcium Leak Drives the Acute Cardiovascular Manifestations of Ciguatera Poisoning: implications for an emerging climate-sensitive cardiotoxic disease

**DOI:** 10.64898/2026.08.31.748269

**Authors:** Hugo Benoit, Ismael Choukri, Pierre Sicard, Elisa Bounasri, Gilles Labesse, Laurie Alburquerque, Mohamed Laabir, Jacques Mercier, Jérôme Thireau, Alain Lacampagne, Albano C. Meli

## Abstract

**Background:** Ciguatera is the most prevalent non-bacterial seafood poisoning worldwide. It results from the consumption of fish contaminated with ciguatoxins (CTXs). Cardiovascular manifestations are frequently reported in affected patients; however, the molecular mechanisms underlying CTX-induced cardiac dysfunction remain poorly understood.

**Methods:** We investigated the effects of purified CTX-1B and CTX-3C on human induced pluripotent stem-cell-derived ventricular cardiomyocytes. Electrophysiology, intracellular calcium cycling and contractility, and cellular structure were assessed. The RyR2 channel activity was studied in planar lipid bilayers, and molecular docking was used to identify a potential CTX-1B interaction site. Consequences on cardiac function were assessed in rats using echocardiography and EKG.

**Findings:** We found that ciguatoxin exposure rapidly increased diastolic intracellular calcium, with severe impacts on the cardiac excitation-contraction coupling, and caused sarcomeric disorganization in human cardiomyocytes. Single-channel recordings indicated that CTX-1B prolonged RyR2 openings at resting cytosolic calcium and altered ATP-dependent gating. These findings are consistent with a state-dependent remodeling of RyR2 gating rather than uniform potentiation. Docking placed CTX-1B within the RyR2 transmembrane region, near ATP-associated regulatory elements. In rats, CTX-1B caused premature atrial contractions, ventricular ectopy, and acute cardiac dysfunction, with impaired relaxation.

**Interpretation:** These findings identify RyR2-dependent calcium leak as a plausible contributor to ciguatoxin cardiotoxicity and extend the established model based on sodium-channel activation and altered sarcolemmal ion handling. Combined disruption of membrane excitability and intracellular calcium release could explain the rapid electrical and mechanical abnormalities associated with ciguatera. As intensive international shipping traffic and the global dispersal of harmful microalgae via ballast water expand exposure risk alongside the international seafood trade, ciguatera should be considered an emerging global cardiovascular hazard.

**Funding:** This work was supported by grants of MUSE CIBSEEA and Exposum Exaltox, ANR MUSAGE (ANR-21-CE14-0026) and FENICE (ANR-22-CE17-0012) and the “Institut National pour la Santé et la Recherche Médicale” (INSERM).

## Introduction

Ciguatera is the most common form of non-bacterial seafood poisoning and results from the consumption of fish containing ciguatoxins (CTXs), potent marine toxins that are accumulated by fish through the food web. Ciguatoxins are potent polyether compounds produced by benthic dinoflagellates belonging primarily to the genera Gambierdiscus and Fukuyoa (1). Considering the global burden of marine biotoxin poisoning, it is clear that these intoxications represent a significant public health challenge. Recent findings on paralytic shellfish poisoning indicate that such events occur regularly across all continents, highlighting persistent gaps in epidemiological surveillance and case reporting (2). In coral reef ecosystems, ciguatoxins enter the food web and accumulate in both herbivorous and carnivorous fish, ultimately posing a risk to human health through the consumption of contaminated fish and causing ciguatera fish poisoning.

Historically, ciguatera has been primarily associated with tropical and subtropical coral reef ecosystems. However, global warming and rising sea surface temperatures may favor the geographical expansion of ciguatera, with cases increasingly reported at higher latitudes, particularly in Atlantic islands and, more recently, in the Mediterranean region (3). More specifically, the degradation of coral reefs, the intensification of maritime transport, and the increasing globalization of fisheries may contribute to the geographical expansion of ciguatera exposure. Ciguatera is characterized by a range of gastrointestinal, neurological, and cardiovascular symptoms. Cardiovascular manifestations include bradycardia, hypotension, and cardiac arrhythmias, highlighting the susceptibility of the cardiovascular system to ciguatoxin-induced toxicity (4). Studies have shown that CTXs are able to bind to voltage-gated sodium channels leading to membrane depolarization and, consequently, secondary alterations in potassium and calcium (Ca^2+^ ) channel activity (5, 6). Although altered Na handling is likely to contribute to CTX-induced cardiac dysfunction, it is unlikely to be the sole mechanism involved. The highly lipophilic nature of CTXs raises the possibility of interactions with intracellular membrane-associated proteins involved in Ca^2+^ handling, including RyR2, a key regulator of sarcoplasmic reticulum Ca^2+^ release and cardiac excitation-contraction coupling. Indeed, the intracellular cardiac ryanodine receptor (RyR2) releases Ca^2+^ from the sarcoplasmic reticulum during excitation-contraction coupling. It is the major actor to increase intracellular Ca^2+^ concentration to allow contraction (*i.e.*, systole). Abnormal RyR2 opening raises diastolic Ca^2+^, reduces the Ca^2+^ available for systolic release, and promotes delayed afterdepolarisations and contractile dysfunction (7). We therefore tested whether RyR2 contributes to CTX cardiotoxicity. Purified CTX-1B and CTX-3C were investigated in human induced pluripotent stem-cell-derived cardiomyocytes (hiPSC-CMs), porcine RyR2 channels incorporated into planar lipid bilayers, and also *in vivo* in rats. We combined intracellular Ca^2+^ imaging, cellular morphology, contractility, multielectrode-array recordings, electrocardiography, echocardiography, and molecular docking to define how CTXs alter intracellular Ca^2+^ regulation and cardiac function.

## Methods

### Study models and toxin exposure

The control hiPSC line AG08C5 (RRID: CVCL_D0KP) was differentiated into ventricular-like cardiomyocytes using temporal modulation of Wnt signalling, followed by lactate selection and maturation to day 30, as described previously (8). Dissociated hiPSC-CMs and two-dimensional cardiac sheets were exposed to vehicle, CTX-1B, or CTX-3C (5 nM) for 5 min or 120 min. The study was conducted in accordance with the Declaration of Helsinki, with written informed consent for hiPSC generation. Animal procedures complied with Directive 2010/63/EU and were approved locally (APAFIS#46232-2023120809257249 v5). Male Wistar rats received intravenous vehicle or CTX-1B (0.13 ng/g bodyweight).

### Ca^2+^ imaging, morphology, electrophysiology, and contraction

Diastolic cytosolic Ca^2+^ was measured with Indo-1 AM in Tyrode solution. Spontaneous Ca^2+^ transients were recorded at 37°C with Fluo-4 AM by confocal line scanning and analysed for frequency, amplitude, decay, and upstroke velocity (9). For morphology, dissociated cells were labelled for alpha-actinin, cardiac troponin I, and DAPI; confocal z-stacks were analysed with ImageJ and MorphoScript (10). Spontaneous contractions of cardiac sheets were recorded for 25 s and analysed from pixel displacement to obtain beat rate, contraction amplitude, and resting time (8). Field potentials were recorded from cardiac sheets with a 64-electrode array before and after 5 nM CTX-1B.

### RyR2 single-channel recordings and docking

Sarcoplasmic-reticulum microsomes from porcine left ventricle were incorporated into planar lipid bilayers. The cis chamber represented the cytoplasmic side and contained defined free Ca^2+^ concentrations calculated with WinMaxC; the trans chamber represented the luminal side. Recordings were obtained at 0 mV and 23°C, filtered at 1 kHz, and digitised at 4 kHz as previously reported (11). Acute experiments were performed at 150 nM cis Ca^2+^ with CTX-1B (1 nM), Na-ATP (25 microM), or sequential addition of ATP then CTX-1B. Two-minute stabilised segments were analysed for open probability (Po), mean open time (To), mean closed time (Tc), and opening frequency (Fo). RyR2 identity was verified by ryanodine-sensitive behaviour. CTX-1B was docked to the open RyR2 cryo-EM structure (PDB 7UA4) using PLANTS, with the search centred on F4853 (12).

#### *In-vivo* assessment

Continuous lead-II electrocardiograms were acquired by jacketed telemetry and analysed for RR and corrected QT intervals and arrhythmic events (13). High-resolution transthoracic echocardiography was performed using a Vevo F2 system with a UHF22x probe (VisualSonics/FUJIFILM). Rats were anaesthetised with 2% isoflurane, maintained at 37°C, and continuously monitored by ECG. B- and M-mode imaging was used to measure ejection fraction and fractional shortening, while radial, longitudinal, and circumferential strain were analysed using Vevo LAB 2.0. Diastolic function was assessed by pulsed-wave and tissue Doppler measurements of E, IVRT, e′, and longitudinal early diastolic strain rate (SRe). Measurements followed established guidelines for echocardiography in rats (14). Experimental details and sample sizes are given in the figure legends.

### Statistical analysis

Normality was assessed with the Shapiro-Wilk test. Two independent groups were compared with an unpaired t test or Mann-Whitney test, and three groups with one-way ANOVA or Kruskal-Wallis analysis, as appropriate. Repeated measurements were analysed by repeated-measures ANOVA with Dunnett correction. Categorical outcomes were compared with Fisher exact tests. Data are shown as individual observations with mean and SEM. Tests were two-sided, and p<0.05 was considered significant. Analyses were done in GraphPad Prism 10.

## Results

### Ciguatoxins rapidly and persistently elevate diastolic intracellular Ca^2+^ and disrupt ECC and Ca^2+^ transients in hiPSC-CMs

Previous study indicated that CTX acts directly on cardiac sodium channels, causing sodium ions influx and reducing the effectiveness of the Na^+^/Ca^2+^ exchanger in guinea-pig cardiac muscle. By this mechanism, CTXs are expected to increase intracellular SR Ca^2+^ release and enhance contractility (15). To determine whether CTXs affect intracellular Ca^2+^ homeostasis in human cardiomyocytes, we first quantified diastolic cytosolic Ca^2+^ levels in ventricular-like healthy control hiPSC-CMs. Acute exposure to either CTX-1B or CTX-3C (5 nM) caused a rapid increase in resting Ca^2+^ detectable within 5 min (Fig. 1A-B). Compared with vehicle, CTX-1B elevated diastolic Ca^2+^ by approximately ∼34.40%, and this elevation remained stable after 2 h of exposure, indicating persistent Ca^2+^ dysregulation. Dose-response experiments revealed that very low concentrations of CTX-1B and CTX-3C (0.3-5 nM, 5 min exposure) were sufficient to significantly increase resting Ca^2+^ (Fig. 1C-D). Notably, the Ca^2+^ elevation reached near-maximal levels at 0.3 nM, with no additional increase at higher concentrations, indicating high potency. The rapid rise in diastolic Ca^2+^ occurred within the first minutes of exposure and remained elevated throughout the entire recording period (Fig. 1E), confirming that CTX induces a sustained disturbance of intracellular Ca^2+^ homeostasis.

**Figure 1.**
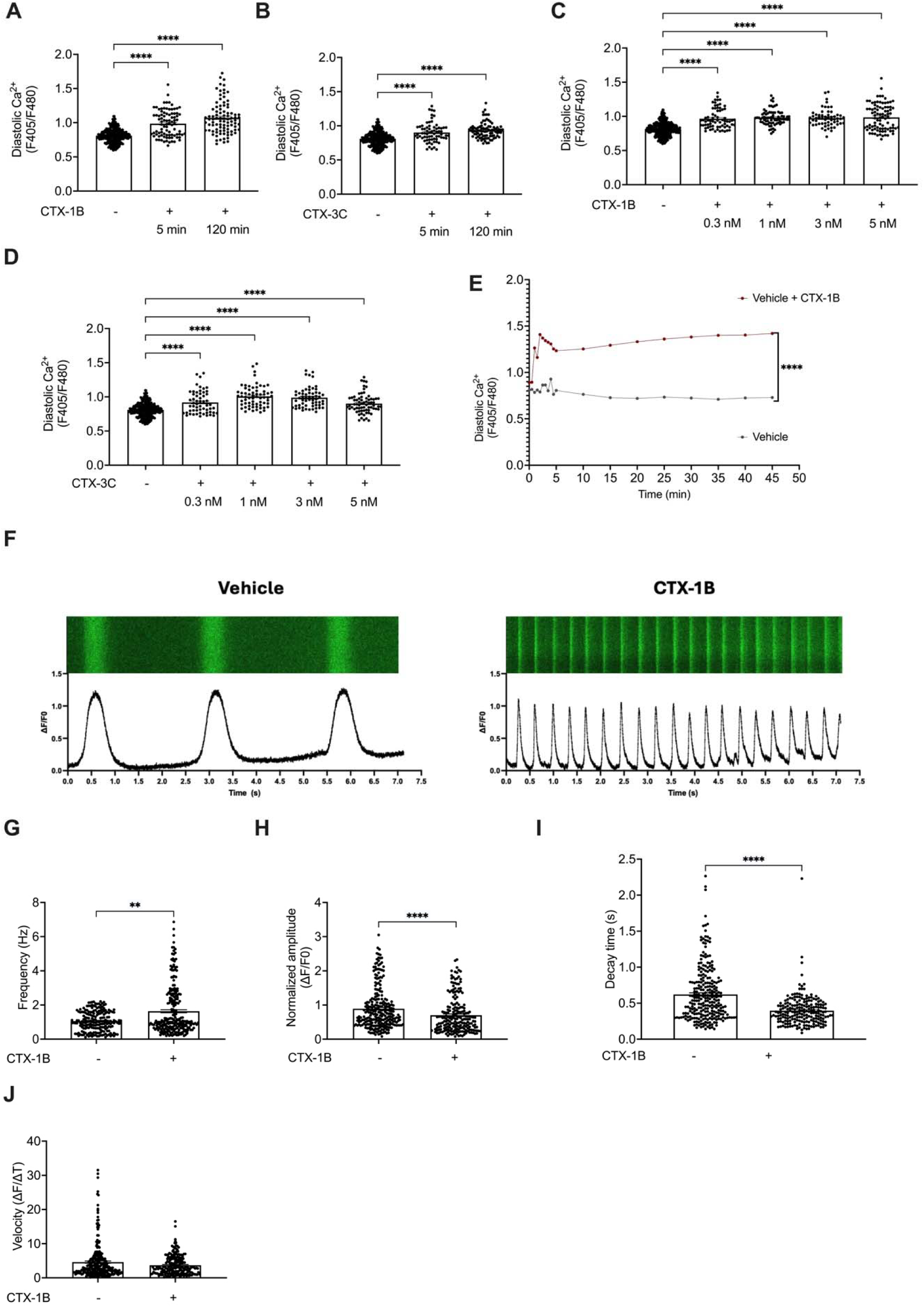
CTX-1B rapidly and persistently elevates diastolic Ca^2+^ levels and altered spontaneous Ca^2+^ transients in hiPSC-CMs. (A-B) Diastolic Ca^2+^ measured after 5 and 120 min of exposure to 5 nM CTX-1B (A) or 5 nM CTX-3C (B) or vehicle in hiPSC-CMs. CTX-1B induced a rapid and sustained increase in resting intracellular Ca^2+^ . (C-D) Dose–response analysis showing diastolic Ca^2+^ levels after 5 min exposure to increasing concentrations of CTX-1B (0.3, 1, and 3 nM) (C) or CTX-3C (D). (E) Representative continuous recording of resting Ca^2+^ levels in hiPSC-CMs exposed to CTX-1B (5 nM, red) or vehicle (in grey). CTX-1B induced a rapid increase in diastolic Ca^2+^ within the first minutes, followed by a persistent elevation throughout the recording period, while control cells displayed stable or slightly decreasing Ca^2+^ levels over time. (F) Representation of spontaneous Ca^2+^ transients in hiPSC-CMs treated with vehicle (left panel) or 5 nM of CTX-1B (right panel). Data were recorded in line-scan configuration using Fluo4-AM with temporal profiles displayed below. (G) Scatter plot showing the Ca^2+^ transient frequency (Hz) in hiPSC-CMs treated with with vehicle or 5 nM of CTX-1B for 2 hours at 37 °C. (H) Scatter plot showing the Ca^2+^ transient normalized amplitude (ΔF/F0) in hiPSC-CMs treated with vehicle or 5 nM of CTX-1B. (I) Scatter plot showing the Ca^2+^ transient decay time (s) in hiPSC-CMs treated with vehicle or 5 nM of CTX-1B. (J) Scatter plot showing the Ca^2+^ transient velocity (ΔF/ΔT) in hiPSC-CMs treated with vehicle or 5 nM of CTX-1B. Data are presented as individual points with mean ± SEM.

Line-scan confocal Ca^2+^ imaging was performed in hiPSC-CMs using the fluorescent Ca^2+^ indicator Fluo-4 AM. Exposure to CTX-1B (5 nM) profoundly distorted the morphology of Ca^2+^ transients (Fig. 1F). Untreated hiPSC-CMs displayed regular and spontaneous Ca^2+^ transients characterized by high amplitude and consistent frequency. In contrast, exposure to 5 nM CTX-1B markedly altered transient morphology (Fig. 1F). CTX significantly increased the frequency of spontaneous Ca^2+^ transients (+54.47% increase) and reduced the transient amplitude by ∼21.22% (Fig. 1G-H). It increased the decay kinetics but did not affect the release velocity (Fig. 1I and J). Similar effects were observed with CTX-3C, notably with increased frequency and reduced amplitude (Fig. S1A-B). However, CTX-3C reduced the release velocity but did not change the decay kinetics (Fig. S1C-D). These alterations indicate impaired SR Ca^2+^ reuptake, reduced systolic Ca^2+^ release, and diastolic Ca^2+^ instability.

### CTX-1B induces a RyR2 gain-of-function consistent with pathological Ca^2+^ leak

To investigate whether CTX-1B-induced elevation of resting intracellular Ca^2+^ could arise from direct modulation of RyR2, single-channel activity was examined using planar lipid bilayers. Under resting cytosolic Ca^2+^ conditions (150 nM), acute application of CTX-1B (1 nM) significantly increased the mean open time of RyR2 (Fig. 2A-D). Temporal analysis revealed discrete episodes of prolonged channel opening after CTX-1B application, whereas vehicle-treated channels predominantly exhibited brief openings (Fig. 2B). In contrast, the population-level increase in open probability did not reach statistical significance (Fig. 2E), indicating that the most robust acute effect of CTX-1B was the prolongation of individual open events rather than a uniform increase in channel activity across all recordings. Analysis of single-channel current amplitudes showed that CTX-1B altered the relative distribution of opening amplitudes without providing clear evidence for a new conductance state (Fig. 2C).

**Figure 2.**
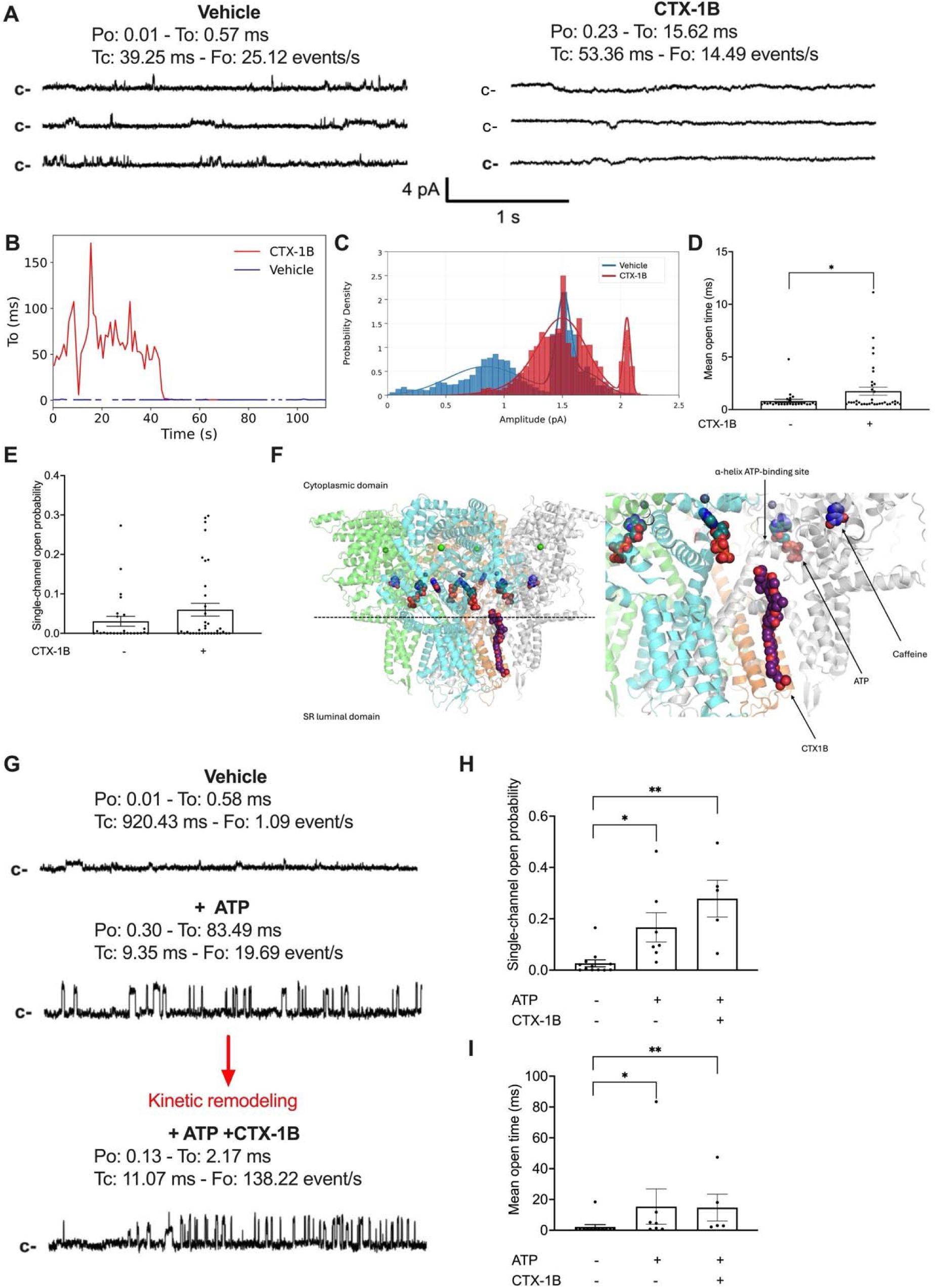
Acute CTX-1B exposure alters RyR2 gating at resting cytosolic Ca^2+^ and modulates ATP-dependent channel activation. (A) Representative single-channel recordings of porcine RyR2 incorporated into planar lipid bilayers under vehicle conditions and following acute exposure to CTX-1B (1 nM) at 150 nM cis Ca^2+^ . Three consecutive segments from the same representative recording are shown for each condition. The closed-channel level is indicated by c-. Open probability (Po), mean open time (To), mean closed time (Tc), and opening frequency (Fo) are reported above the traces. The vertical and horizontal scale bars represent 4 pA and 1 s, respectively. (B) Temporal distribution of open-event durations under vehicle and CTX-1B conditions in representative recordings. CTX-1B produced discrete episodes of prolonged openings, consistent with the transient recruitment of a long-open gating mode. (C) Probability-density distributions of single-channel current amplitudes under vehicle and CTX-1B conditions. Histograms and fitted density curves illustrate the distribution of conductance states detected in the representative recordings. (D, E) Quantification of mean open time (D) and single-channel open probability (E) under vehicle and acute CTX-1B exposure. CTX-1B significantly increased mean open time, whereas the population-level increase in Po did not reach statistical significance. Individual points represent independent channel recordings; bars and error bars represent mean ± SEM. (F) Molecular docking model showing the predicted position of CTX-1B within the RyR2 transmembrane region. The overview indicates the position of the predicted CTX-1B-binding region relative to the cytoplasmic and SR-luminal domains. The enlarged view shows the spatial relationship between docked CTX-1B and the ATP- and caffeine-associated regulatory regions, including the α-helical ATP-binding region. The docking model indicates spatial proximity and provides a structural hypothesis for functional coupling but does not establish direct competition between CTX-1B and ATP. (G) Representative sequential single-channel recording at 150 nM cis Ca^2+^ showing RyR2 activity under vehicle conditions, after addition of Na-ATP (25 µM), and after subsequent addition of CTX-1B (1 nM) in the continued presence of ATP. ATP increased Po, prolonged openings, shortened closed intervals, and increased opening frequency relative to vehicle. Subsequent CTX-1B exposure shifted the ATP-activated channel toward a gating pattern characterized by shorter and markedly more frequent openings, illustrating kinetic remodeling rather than a uniform additive increase in all gating parameters. The closed-channel level is indicated by c-. (H, I) Quantification of single-channel open probability (H) and mean open time (I) under vehicle, ATP, and ATP + CTX-1B conditions. ATP and ATP + CTX-1B increased Po and mean open time relative to vehicle. Individual points represent independent channel recordings; bars and error bars represent mean ± SEM.

Because ATP is an important physiological regulator of RyR2, the effect of CTX-1B was subsequently examined in ATP-activated channels. Addition of Na-ATP (25 µM) at 150 nM cis Ca^2+^ significantly increased RyR2 open probability and mean open time relative to vehicle (Fig. 2G-I). Subsequent addition of CTX-1B in the continued presence of ATP maintained the increase in channel activity but markedly altered the representative gating pattern. In particular, the ATP-associated long openings were redistributed toward shorter, substantially more frequent events. Thus, CTX-1B did not produce a uniform additive increase in every gating parameter but instead remodeled the ATP-activated gating state. This kinetic interaction is consistent with functional coupling between CTX-1B-sensitive and ATP-sensitive regulatory mechanisms within RyR2.

Molecular docking identified a predicted CTX-1B-binding region within the RyR2 transmembrane domain, involving helices encompassing residues 4751-4762 and 4840-4864 and located near F4853 (Fig. 2F). The predicted toxin-binding position lies in spatial proximity to the ATP- and caffeine-associated regulatory regions. This structural arrangement provides a plausible basis for CTX-1B-induced stabilization of RyR2 opening and for the observed remodeling of ATP-dependent gating. However, the docking analysis indicates spatial proximity rather than direct competition and does not establish that CTX-1B occupies the ATP-binding pocket. Collectively, the single-channel and docking results support a model in which CTX-1B binds within the RyR2 transmembrane region and allosterically alters the coupling between cytoplasmic ligand regulation and channel opening, thereby favoring prolonged or otherwise dysregulated openings under resting Ca^2+^ conditions.

### CTX exposure leads to sarcomere elongation and disassembly in hiPSC-CMs and impairs contractile performance of hiPSC-derived cardiac sheets

Next, we assayed whether CTXs could alter the morphology of the hiPSC-CMs using the MorphoScript pipeline (10) and 2 main markers of the sarcomeric organization, alpha-actinin and cardiac troponin I (TnI). Exposure to 5nM CTX-1B and CTX-3C for 2 hours induced significant alterations in both cellular dimensions and the organization of the contractile apparatus. Specifically, the mean sarcomere distance, as defined by alpha-actinin Z-disc spacing, increased significantly following CTX-3C treatment. This structural lengthening, representing a 3.50% increase compared to vehicle controls (Fig. 3A-B) while the cell area remained unchanged (Fig. 3C). This structural lengthening was accompanied by specific shifts in fluorescence intensity profiles. The mean protein intensity for alpha-actinin and TnI markers decreased while the mean back intracellular background increased by 31.68% (Fig. 3D-E). These measurements reflect a loss of structural compartmentalization, with contractile proteins redistributing into the diffuse cytosolic space as the myofibrils elongate. Although global shape descriptors such as rectangularity and eccentricity did not reach statistical significance, a clear trend toward cellular rounding was observed (Fig. S2A-B). Furthermore, the non-significant decrease in granularity and protein distribution (Fig. S2C) metrics supported the transition of contractile proteins from organized sarcomeric filaments into a diffuse cytosolic state, providing a quantitative basis for the observed myofibrillar disassembly. Collectively, these morphometric data quantify a rapid reorganization of the cardiomyocyte cytoarchitecture characterized by sarcomeric dilatation and protein disassembly.

**Figure 3.**
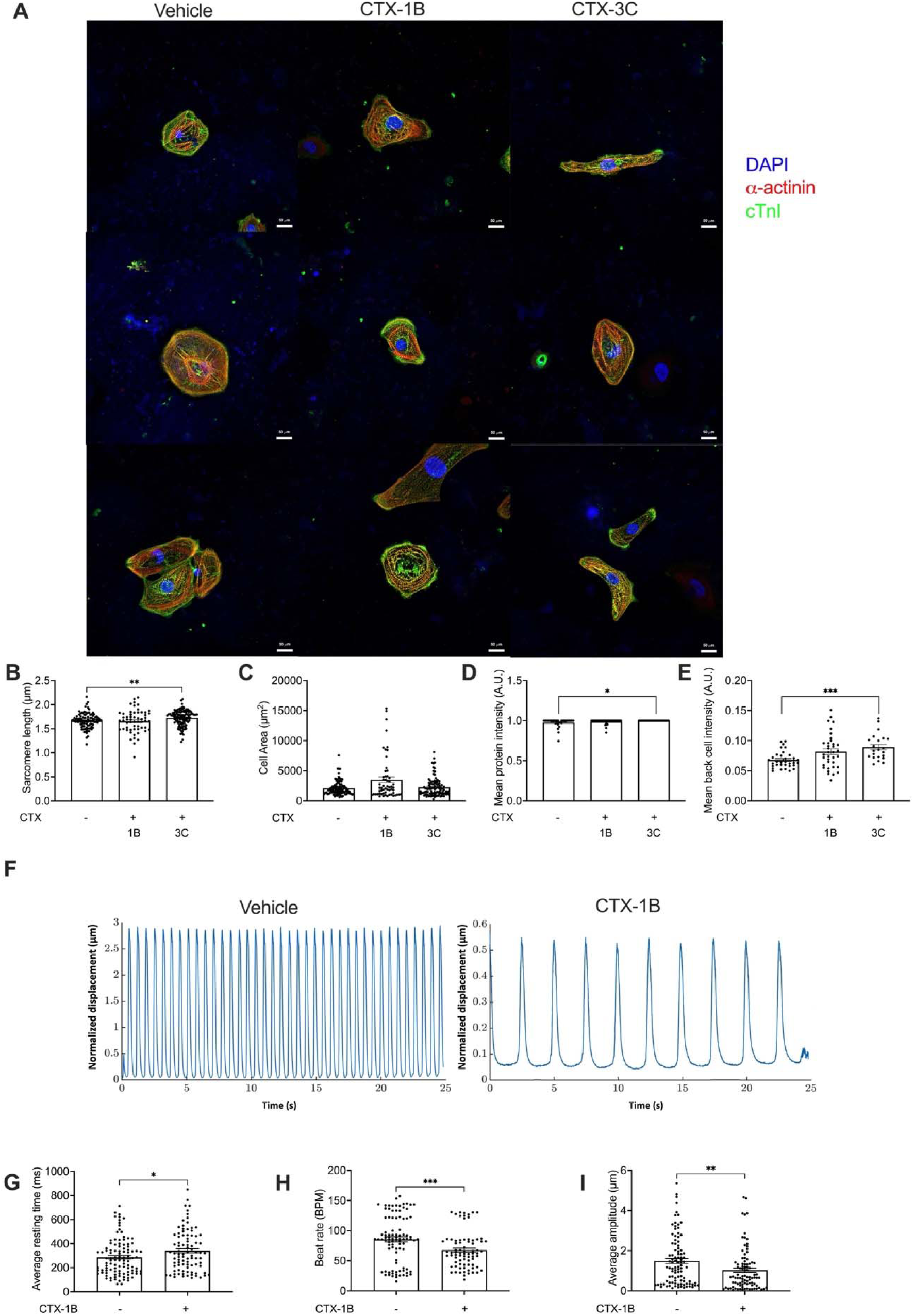
CTX exposure alters sarcomeric organization and contractile properties in hiPSC-derived cardiomyocytes. (A) Representative immunofluorescence images of dissociated hiPSC-CMs treated with vehicle, 5 nM CTX-1B, or 5 nM CTX-3C for 120 min. Cells were labelled for α-actinin (red) and cardiac troponin I (cTnI; green) to assess sarcomeric organization, and nuclei were counterstained with DAPI (blue). Scale bars, 50 µm. (B) Quantification of sarcomere length, determined from α-actinin-positive Z-disc spacing. (C) Quantification of hiPSC-CM surface area. (D) Quantification of mean sarcomeric protein fluorescence intensity. (E) Quantification of mean cellular background fluorescence intensity. (F) Representative displacement traces from spontaneously contracting hiPSC-CM sheets under vehicle conditions or following CTX-1B exposure. (G-I) Quantification of average resting time (G), spontaneous beat rate (H), and average contraction amplitude (I) in vehicle- and 5nM CTX-1B-treated cardiac sheets (for 120 min). Individual points represent cells or analyzed contractions, as appropriate, and bars and error bars indicate mean ± SEM.

We next examined whether the Ca^2+^ and electrophysiological alterations translated into contractile dysfunction. Two-dimensional cardiac sheets of hiPSC-CMs were imaged using high-speed confocal video microscopy and contractile motion was quantified through MATLAB- and vector-based displacement analysis. In the presence of 5 nM CTX-1B, the contractile performance was markedly impaired, with slower resting time and diminished beat frequency (by 20.87%) and contraction amplitude (by 31.15%) compared with untreated controls (Fig. 3F–I). In the presence of 5 nM CTX- 3C, identical contractile defects were accurately observed (Fig. S2D-F). These changes are consistent with weakened Ca^2+^ transients and decreased intracellular Ca^2+^ availability during systole. Functionally, they parallel the bradycardia and myocardial depression observed in clinical cases of ciguatera fish poisoning (16).

### CTX-1B suppresses spontaneous electrical activity and alters field potential duration in cardiac network in vitro and induces bradycardia and arrhythmias in vivo

Using multielectrode array recordings, we next assessed the electrophysiological consequences of CTX exposure at the tissue level using spontaneously beating 2D cardiac sheets. Under baseline conditions, cells exhibited stable and rhythmic field potentials with consistent depolarization and repolarization phases. Treatment with 5 nM CTX-1B or CTX-3C markedly reduced the number of field potentials and shortened field potential duration, while leaving amplitude unaffected (Fig. 4A-B). Quantitatively, CTX-1B exposure decreased the number of spontaneous field potentials per recording period by 66.23% and reduced the duration of individual depolarization events by 41.91%, indicating impaired repolarization dynamics (Fig. 4C-E). The preservation of signal amplitude suggests that conduction integrity remained intact despite altered excitability (Fig. 4D). These electrophysiological findings align with the Ca^2+^ imaging results, indicating that disrupted Ca^2+^ cycling and RyR2-mediated Ca^2+^ leak compromise automaticity and electrical stability, potentially predisposing cells to arrhythmic behavior.

**Figure 4.**
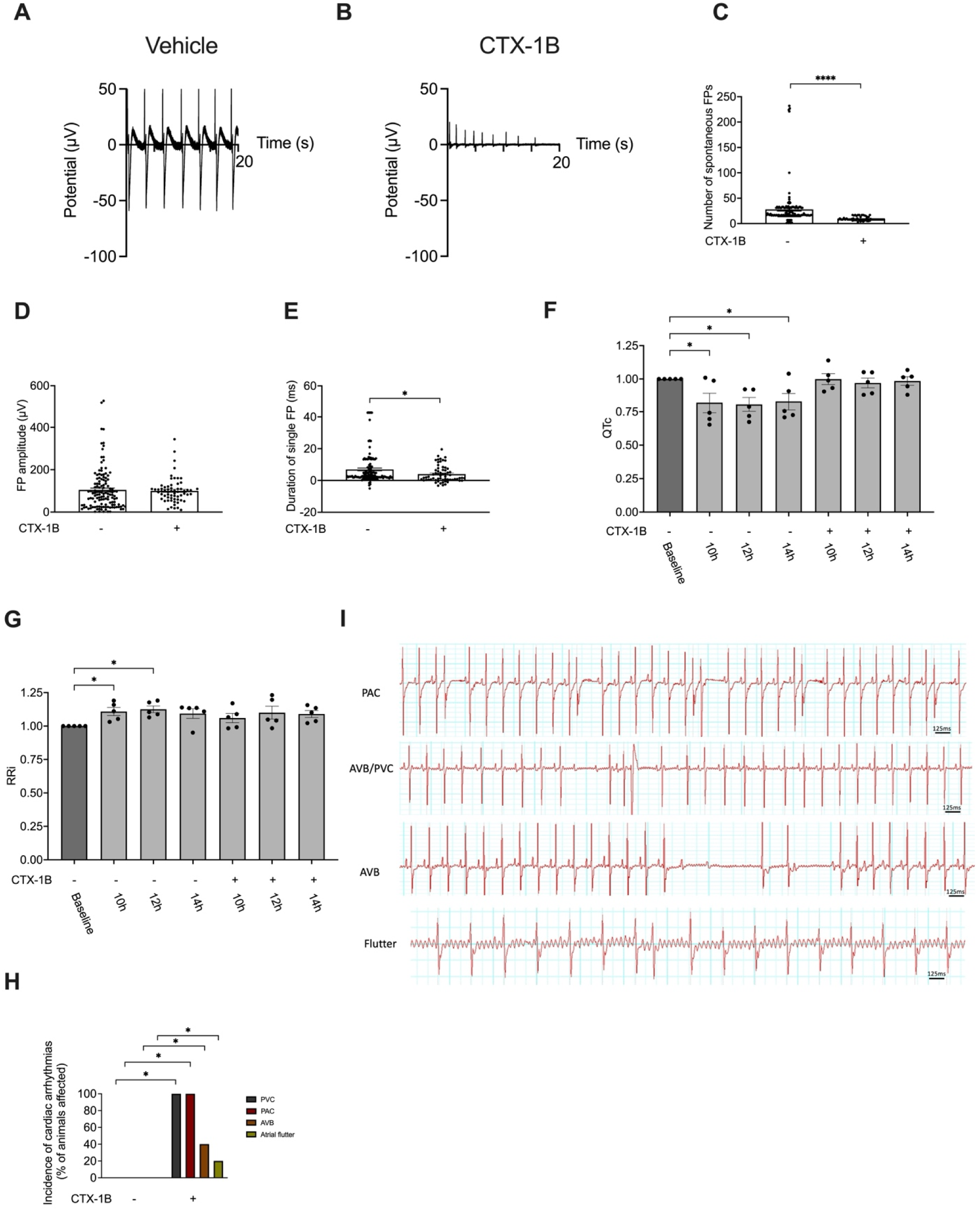
CTX-1B disrupts spontaneous field potentials in hiPSC-CMs and causes cardiac arrhythmias and conduction defects in anaesthetized rats. (A) Representative MEA trace of spontaneous field potentials (FPs) recorded under control conditions, showing regular rhythmic activity. (B) Representative trace after CTX exposure, illustrating suppression of spontaneous activity and shortened FP duration. (C) Quantification of the number of spontaneous FPs per recording, showing a significant reduction following CTX. (D) FP amplitude remained unchanged between control and CTX groups. (E) Duration of single FP events was significantly shortened in the presence of CTX. (F) Corrected QT interval (QTc, Mitchell formula) derived from continuous ECG recordings in conscious, unrestrained male Wistar rats (n=6–8 per group) administered intravenous CTX-1B or vehicle. Recordings spanned baseline (pre-injection) and post-injection timepoints at 10 h, 12 h and 14 h for both CTX-1B-treated (+) and control (−) animals. Values are normalized to individual baseline (pre-injection). (G) Normalized RR interval (RRi, relative to baseline) derived from continuous ECG recordings in conscious, unrestrained male Wistar rats (n=6–8 per group) administered intravenous CTX-1B or vehicle. (* p<0.05 vs. baseline; repeated-measures ANOVA with Dunnett’s post-hoc) at 10 h and 12 h post-injection in CTX-1B-treated rats, indicative of bradycardia, with partial recovery by 14 h. Horizontal bars denote timepoints sampled for CTX-1B (+) and control (−) groups. (H) Representative ECG recordings illustrating rhythm and conduction abnormalities observed following CTX-1B exposure, including premature atrial contractions (PACs), atrioventricular block associated with premature ventricular contractions (AVB/PVCs), atrioventricular block (AVB), and atrial flutter. Horizontal scale bars represent 125 ms. (I) Incidence of cardiac arrhythmias and conduction abnormalities (% of animals affected) during the 10–14 h post-injection window. CTX-1B markedly increased the prevalence of PAC and PVC, with trends towards higher AVB and atrial flutter (* p<0.05, Fisher’s exact test vs. control). Data are presented as individual points with mean ± SEM.

To validate these observations under physiological conditions, we monitored cardiac function in freely moving Wistar rats intravenously injected either with vehicle or with CTX-1B (0.13 ng/g body weight). We observed that CTX induced a significant bradycardic response when compared to baseline whereas vehicle did not. In addition, when compared to post-vehicle injection, CTX lengthened the ventricular repolarization (Fig. 4F-G). Nevertheless, we have to note that difference probably account for the decrease in QTc in vehicle group, according to circadian variation whereas it surprisingly remained unchanged on CTX group. In addition, animals treated with CTX developed cardiac arrhythmias, including atrioventricular block associated with premature ventricular contractions (5/5) whereas control group did not. Atrioventricular blocks were also observed in 2/5 animals (Fig. 4H-I). These data confirm that systemic exposure to CTX-1B induces complex electrophysiological disturbances consistent with Ca^2+^ -handling abnormalities and RyR2 hyperactivation.

### CTX-1B induces impaired contractile function in vivo resulting in an acute heart failure with preserved ejection fraction

We next assessed cardiac function by echocardiography in vehicle- and CTX-1B-treated rats. CTX-1B did not significantly alter heart rate or left ventricular ejection fraction (Fig. 5A-B). Despite preserved ejection fraction, fractional shortening and global radial strain were significantly reduced after CTX-1B exposure (Fig. 5C-D), indicating impaired radial systolic function. Global longitudinal strain was also significantly reduced, whereas global circumferential strain remained unchanged (Fig. 5E-F). CTX-1B additionally impaired myocardial relaxation. Isovolumic relaxation time was significantly prolonged, and the early diastolic strain-rate parameter SRe was significantly reduced in CTX-1B-treated animals (Fig. 5G-H). Consistent with increased left ventricular filling pressure, the E/SRe ratio was significantly elevated (Fig. 5I). Tissue Doppler analysis further showed a significant reduction in the E/e′ parameter, whereas the transmitral E/A ratio was unchanged (Fig. 5J-K). Taken together, these results indicate that acute CTX-1B exposure leads to both systolic and diastolic dysfunction despite preserved global ejection fraction. The reduction in fractional shortening and radial and longitudinal strain demonstrated subtle systolic impairment uncaptured by conventional ejection-fraction measurements, whereas prolonged isovolumic relaxation, reduced SRe, and altered diastolic ratios indicate impaired ventricular relaxation. These results support an acute cardiac dysfunction phenotype with preserved ejection fraction, rather than establishing clinical heart failure with preserved ejection fraction.

**Figure 5.**
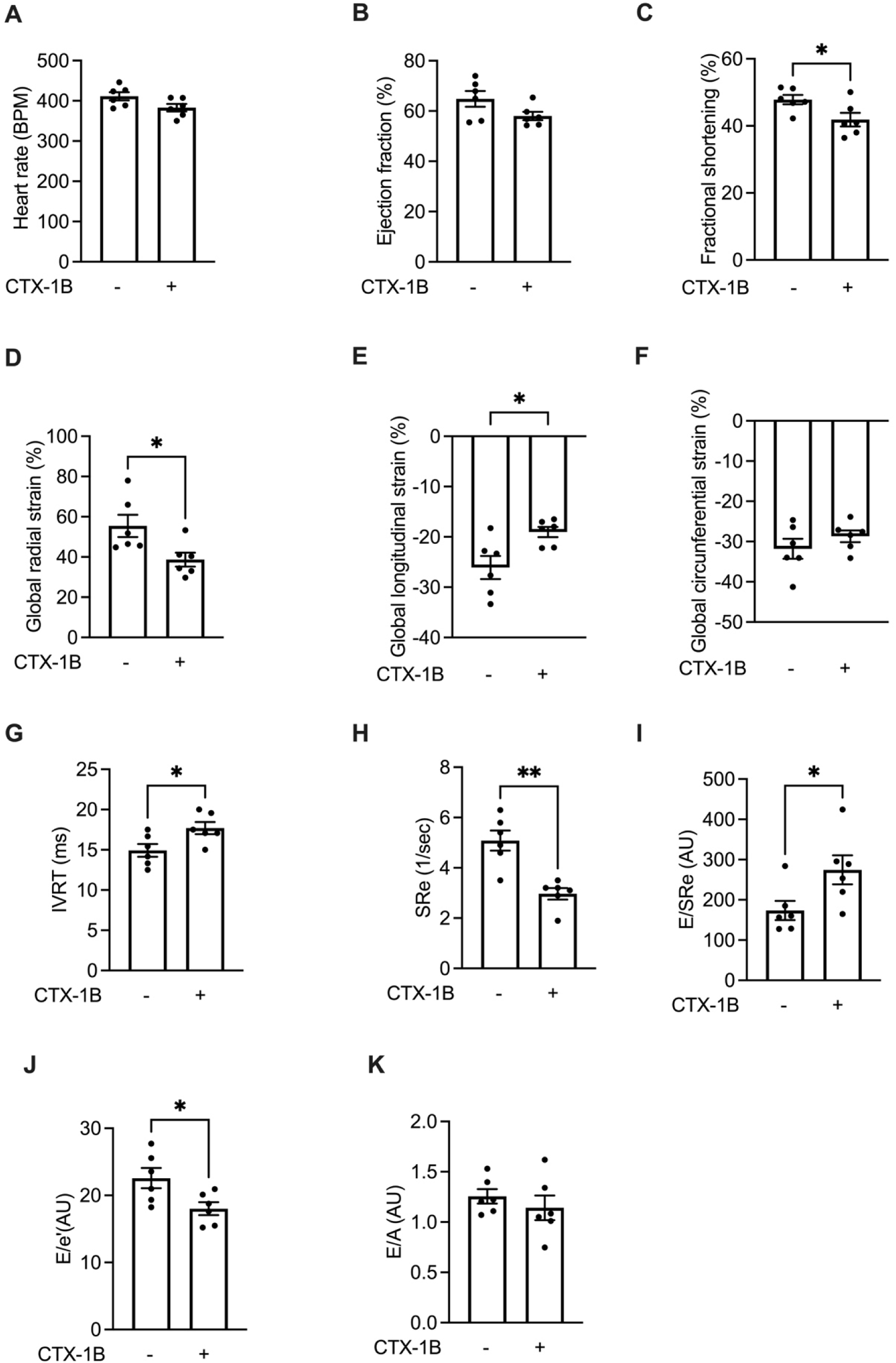
Acute CTX-1B exposure impairs systolic and diastolic cardiac function despite preserved ejection fraction in rats. Cardiac function was assessed by echocardiography following intravenous administration of vehicle or CTX-1B (0.13 ng/g bodyweight). (A) Heart rate. (B) Left ventricular ejection fraction. (C) Fractional shortening. (D-F) Global radial (D), longitudinal (E), and circumferential (F) strain. (G) Isovolumic relaxation time (IVRT). (H) Early diastolic strain rate (SRe). (I) Ratio of early transmitral flow velocity to early diastolic strain rate (E/SRe). (J) Ratio of early transmitral flow velocity to early diastolic mitral annular velocity (E/e′). (K) Ratio of early to late transmitral flow velocity (E/A). CTX-1B significantly reduced fractional shortening and global radial and longitudinal strain, whereas ejection fraction and global circumferential strain remained unchanged. CTX-1B also prolonged IVRT, reduced SRe, increased E/SRe, and reduced E/e′, while the transmitral E/A ratio was unchanged. These findings indicate combined systolic and diastolic dysfunction that was not detected by conventional measurement of ejection fraction. Data are presented as individual observations with mean ± SEM.

## Discussion

Ciguatera is an increasingly important marine biotoxin-related foodborne illness, yet the mechanisms underlying its cardiovascular manifestations remain poorly understood (17, 18). Our findings extend the established model of persistent voltage-gated sodium-channel activation by showing that CTXs also disrupt intracellular Ca^2+^ handling. In human cardiomyocytes, CTX exposure rapidly and persistently increased diastolic Ca^2+^ , altered spontaneous Ca^2+^ transients, impaired contraction, and disrupted sarcomeric organisation. In parallel, CTX-1B modified RyR2 gating in planar lipid bilayers and caused electrical and mechanical cardiac dysfunction in rats.

## CTXs promote RyR2-dependent Ca^2+^ leak

CTX-dependent Na entry can increase intracellular Ca^2+^ through altered Na^+^/Ca^2+^ exchanger activity (15). Our data identify RyR2 modulation as a complementary mechanism. At 150 nM cytosolic Ca^2+^ , acute CTX-1B prolonged individual RyR2 openings without clear evidence of a new conductance state. Prolonged openings would increase time-integrated sarcoplasmic-reticulum Ca^2+^ efflux, even though the population increase in open probability did not reach significance. This mechanism provides a plausible explanation for the rapid elevation of diastolic Ca^2+^ and resembles aspects of pathological RyR2 leak reported in inherited arrhythmogenic disorders and heart failure (8, 19, 20).

The ATP experiments further suggest that CTX-1B is not simply an independent RyR2 agonist. ATP increased RyR2 open probability and mean open time, whereas subsequent CTX-1B shifted the representative gating pattern towards shorter, more frequent openings. The combined condition therefore reflected kinetic redistribution rather than uniform additive or synergistic activation. Molecular docking placed CTX-1B within the RyR2 transmembrane region near F4853 and close to ATP- and caffeine-associated regulatory elements. Together, these findings support state-dependent coupling between CTX-sensitive and ATP-sensitive gating pathways. They do not, however, demonstrate that CTX-1B occupies or competes for the ATP-binding pocket. Structural studies and targeted mutagenesis will be needed to distinguish local steric effects from longer-range allosteric communication.

### Consequences for excitation-contraction coupling and cellular structure

CTX exposure increased the frequency of spontaneous Ca^2+^ transients while reducing their amplitude and slowing Ca^2+^ removal. This pattern is compatible with inefficient Ca^2+^ cycling, in which diastolic sarcoplasmic-reticulum leak limits the Ca^2+^ available for systolic release and relaxation. The accompanying reduction in contraction amplitude and prolongation of resting time support this interpretation and resemble Ca^2+^ -handling abnormalities described in failing myocardium (21).

Ca^2+^ dysregulation was also associated with rapid structural changes. CTX-3C increased sarcomere length, reduced the organization of α-actinin and cardiac troponin I signals, and increased diffuse intracellular fluorescence. These findings are consistent with reduced compartmentalization of contractile proteins, although they do not establish proteolysis or protein release into the cytosol. Ca^2+^ -dependent proteases and cytoskeletal remodeling are possible contributors, but this mechanism requires direct testing. The development of structural and mechanical abnormalities within 2 h is nevertheless consistent with the acute onset of cardiac symptoms during severe intoxication.

### Electrical and cardiac consequences and clinical relevance

In cardiac sheets, CTX exposure reduced beat rate, contraction amplitude, and spontaneous field-potential activity. Persistent sodium-channel activation could depolarize the membrane and reduce channel availability, whereas abnormal RyR2-mediated Ca^2+^ release could activate electrogenic Na^+^/Ca^2+^ exchange (11). Depending on timing and magnitude, the resulting currents could suppress automaticity through sustained depolarization or promote delayed afterdepolarizations and ectopic activity.

The *in vivo* observations strongly support the translational relevance of these cellular findings. CTX-treated animals developed bradycardia, premature atrial contractions, ventricular ectopy, and atrioventricular conduction abnormalities, reproducing several cardiovascular manifestations reported in human ciguatera poisoning. While previous studies have mainly attributed these manifestations to autonomic nervous system dysfunction, our results suggest that intrinsic cardiac mechanisms contribute significantly. The identification of RyR2-mediated Ca^2+^ leak provides a mechanistic link between toxin exposure and arrhythmogenesis. Elevated diastolic Ca^2+^ is a well-established trigger of delayed afterdepolarizations and ectopic activity, providing a plausible explanation for the atrial and ventricular arrhythmias observed in vivo. This framework reconciles reduced spontaneous activity *in vitro* with bradycardia, premature contractions, ectopy, and conduction abnormalities *in vivo* (22, 23). Perhaps the most clinically relevant finding is the demonstration that CTX exposure produces an acute form of cardiac dysfunction characterized by preserved ejection fraction despite measurable impairment of myocardial performance. Conventional echocardiographic indices such as ejection fraction remained unchanged, whereas more sensitive parameters including fractional shortening, longitudinal strain, radial strain, IVRT, E/E′, and E′/A′ ratios revealed both systolic and diastolic impairment. This profile closely resembles the early stages of HFpEF, where myocardial mechanics and relaxation are compromised despite preserved global pump function. The combination of RyR2 leak, elevated diastolic Ca^2+^ , impaired relaxation, and increased filling pressures provides a coherent mechanistic framework for these observations as previously reported (11). Notably, the preservation of circumferential strain suggests that longitudinal and radial myocardial functions may be more vulnerable to acute Ca^2+^ dysregulation than circumferential mechanics. Our findings also identify potential therapeutic directions that warrant experimental testing. Flecainide and ranolazine warrant preclinical testing as potential treatments for CTX cardiotoxicity. Flecainide could limit Na 1.5 activity and RyR2-dependent Ca^2+^ release, whereas ranolazine could reduce late Na influx and secondary Ca^2+^ overload. Neither drug has been validated for ciguatera and both require testing in cellular and animal models before clinical consideration.

To synthesize these molecular and clinical findings within a broader planetary health context, we have developed a schematic model illustrating the cardiotoxic pathway of Ciguatera (Fig. S3). While Ciguatera has historically been confined to endemic reef ecosystems, modern anthropogenic pressures are rapidly altering its geographic distribution. Global warming and the associated increase in seawater temperatures, together with intensive international shipping lane traffic and the subsequent discharge of ballast water serve as major vectors for the global dispersal of harmful benthic microalgae, including *Gambierdiscus* species. When coupled with the expanding global seafood trade, these dispersal mechanisms introduce contaminated fish into non-endemic regions. Consequently, the CTX-induced RyR2 dysfunction and resulting cardiovascular abnormalities detailed in this study must now be managed as an emerging global health challenge rather than a localized tropical disease.

### Limitations

Several limitations should be considered. hiPSC-CMs are less mature than adult cardiomyocytes and were derived from one control line. Bilayer recordings isolate RyR2 from much of its native regulatory environment, and docking predicts proximity rather than binding or competition. Sample sizes were modest, and the animal dose was derived from toxicokinetic studies rather than well-defined human dietary exposure (24). Human exposure thresholds remain uncertain and vary with toxin congener and individual susceptibility (6). Finally, this study examined acute exposure and cannot address repeated exposure, recovery, or long-term remodeling.

## Conclusion

CTXs appear to disrupt cardiac function through convergent effects on membrane excitability and intracellular Ca^2+^ release. RyR2 gating modulation provides a mechanistic link between elevated diastolic Ca^2+^ , impaired excitation-contraction coupling, electrical instability, and myocardial dysfunction with preserved ejection fraction. As ciguatera expands beyond historically endemic regions, cardiovascular involvement could become increasingly relevant, particularly in people with pre-existing abnormalities of Ca^2+^ handling. Clinical studies are now needed to define cardiac risk, recovery, and monitoring requirements after intoxication.

## Author contributions

Conceptualization, J.M., A.L. and A.C.M; Investigation, H.B., I.C., P.S., E.B., I.C., P.S., G.L., J.T., A.L., and A.C.M.; Writing, M.L., A.L. and A.C.M.

## Acknowledgement

We thank the infrastructure ChemBioFrance for managing the virtual docking. We thank the Institut NeuroMyoGene - Pathophysiology and Genetics of Neuron and Muscle (INMG-PGNM) for the healthy control hiPSC line (AG08C5). We thank the staff of PhyMedExp animal care facility, Florie Lopez and the Montpellier animal facilities network (RAM, Biocampus).

## Notes

### Competing Interest Statement

The authors have declared no competing interest.

