## Supplemental figures for "RyR2-Dependent Calcium Leak Drives the Acute Cardiovascular Manifestations of Ciguatera Poisoning: implications for an emerging climate-sensitive cardiotoxic disease"

**Supplemental material**

**Figure S1.** (A) Scatter plot showing the Ca²⁺ transient frequency (Hz) in hiPSC-CMs treated with vehicle or 5 nM of CTX-3C for 2 hours at 37 °C. (B) Scatter plot showing the Ca²⁺ transient normalized amplitude (ΔF/F_0_) in hiPSC-CMs treated with vehicle or 5 nM of CTX- 3C. (C) Scatter plot showing the Ca²⁺ transient decay time (s) in hiPSC-CMs treated with vehicle or 5 nM of CTX-3C. (D) Scatter plot showing the Ca²⁺ transient velocity (ΔF/ΔT) in hiPSC-CMs treated with vehicle or 5 nM of CTX-3C. Data are presented as individual points with mean ± SEM.

**Figure S2.** (A) Measurement of Rectangularity (dimensionless ratio), quantifying the extent to which the cell area fills its minimum bounding box of hiPSC-CMs treated with vehicle or 5 nM CTX-1B for 120 min. (B) Measurement of Eccentricity (dimensionless ratio), indicating the elongation of the cardiomyocyte where values closer to 1 represent more elongated, rod-like structures. (C) Calculation of the Distribution index (dimensionless ratio), representing the spatial arrangement of fluorescent signal across the cell body. (D-F) Scatter plots showing the beat rate (BPM) (D), average contraction amplitude (µm) (E) and average resting time (ms) (F) of the untreated and treated cardiac sheets (5 nM of CTX-3C for 120 min). Data are presented as individual points with mean ± SEM.

**Figure S3.** Schematic overview of CTX-induced cardiotoxicity in the context of the global dispersal of harmful benthic microalgae. Intensifying global pressures and perturbations, such as rising sea temperatures, coral reef degradation, increasing international maritime traffic, and the discharge of ballast water contaminated with microalgae, may facilitate the dispersal and geographical expansion of harmful benthic microalgae, including *Gambierdiscus* species. Associated with expanding global seafood trade and extreme weather events, this dispersal broadens the geographic risk of human exposure to CTX-contaminated fish. At the molecular level, CTX-1B allosterically binds to RyR2. This binding induces kinetic remodeling characterized by an increased mean open time (To), driving a pathological SR Ca²⁺ leak into the cytosol. The resulting intracellular Ca²⁺ dysregulation acts as a trigger for delayed afterdepolarizations (DADs), leading to cellular electrical instability, arrhythmias, bradycardia, and atrioventricular (AV) blocks. Ultimately, this RyR2-dependent dysfunction manifests clinically as acute cardiovascular symptoms, including premature contractions and hypotension, posing significant diagnostic challenges and highlighting ciguatera as an expanding planetary health threat for vulnerable populations worldwide.

**Figure S1**

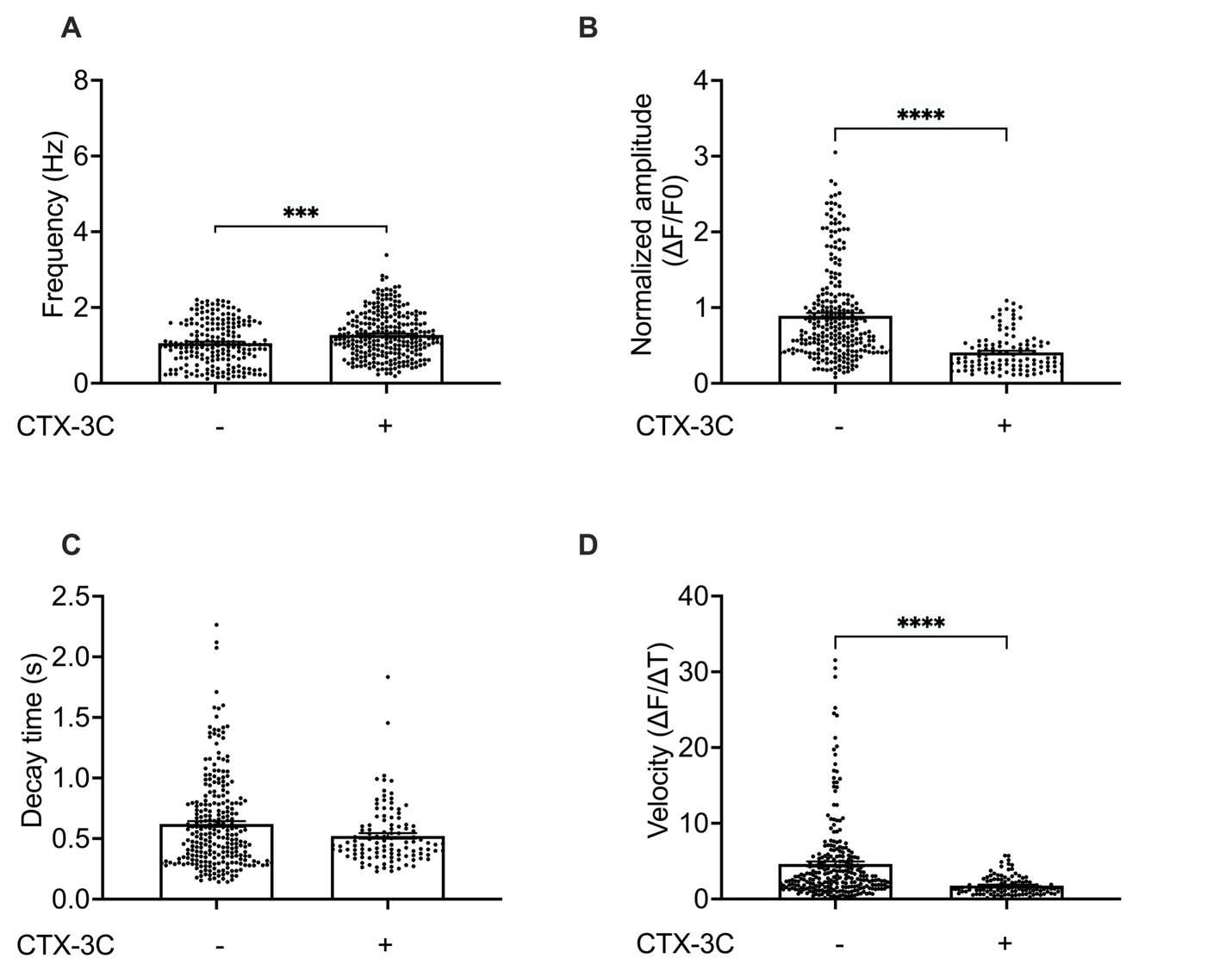

**Figure S2**

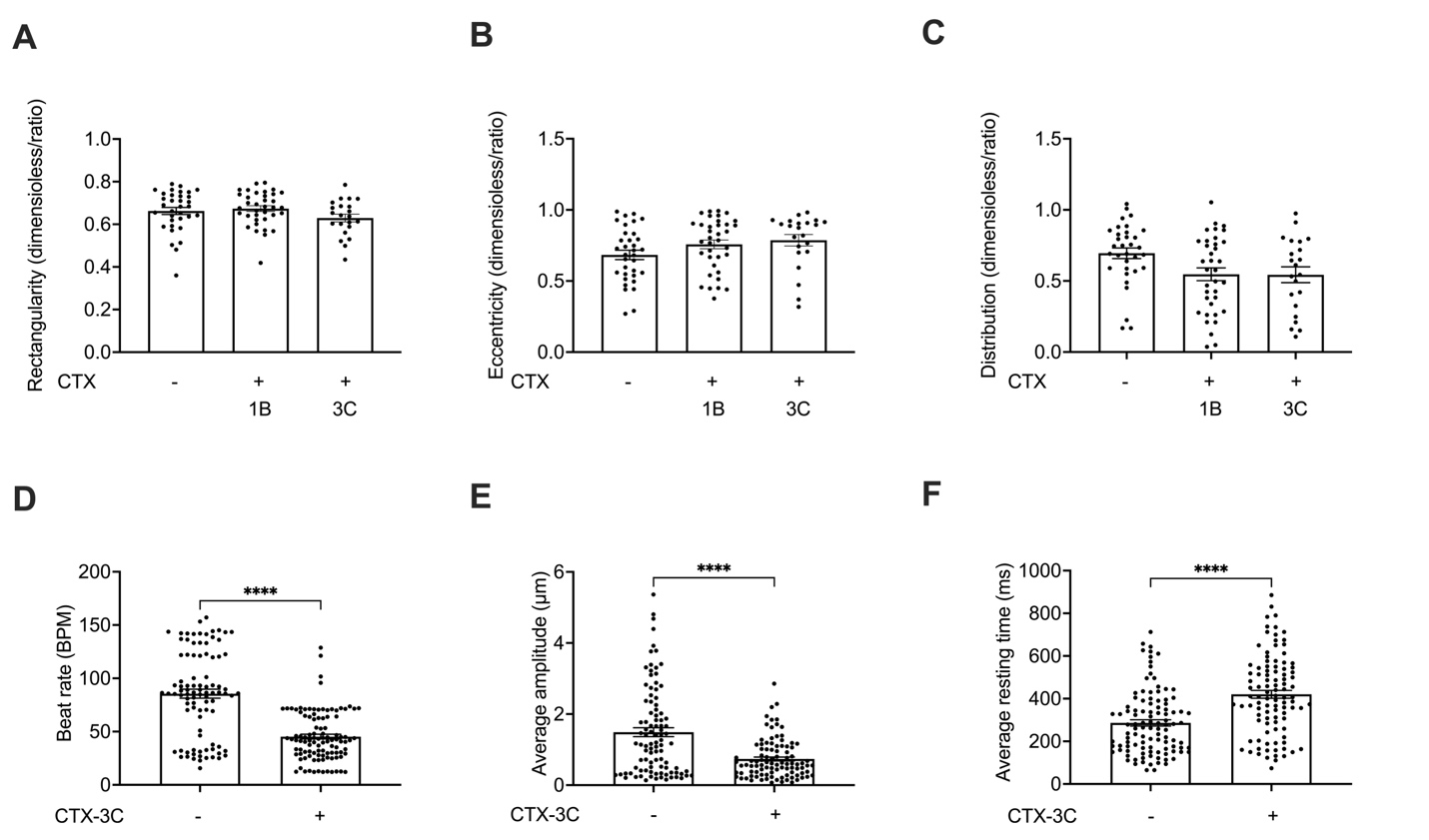

**Figure S3**

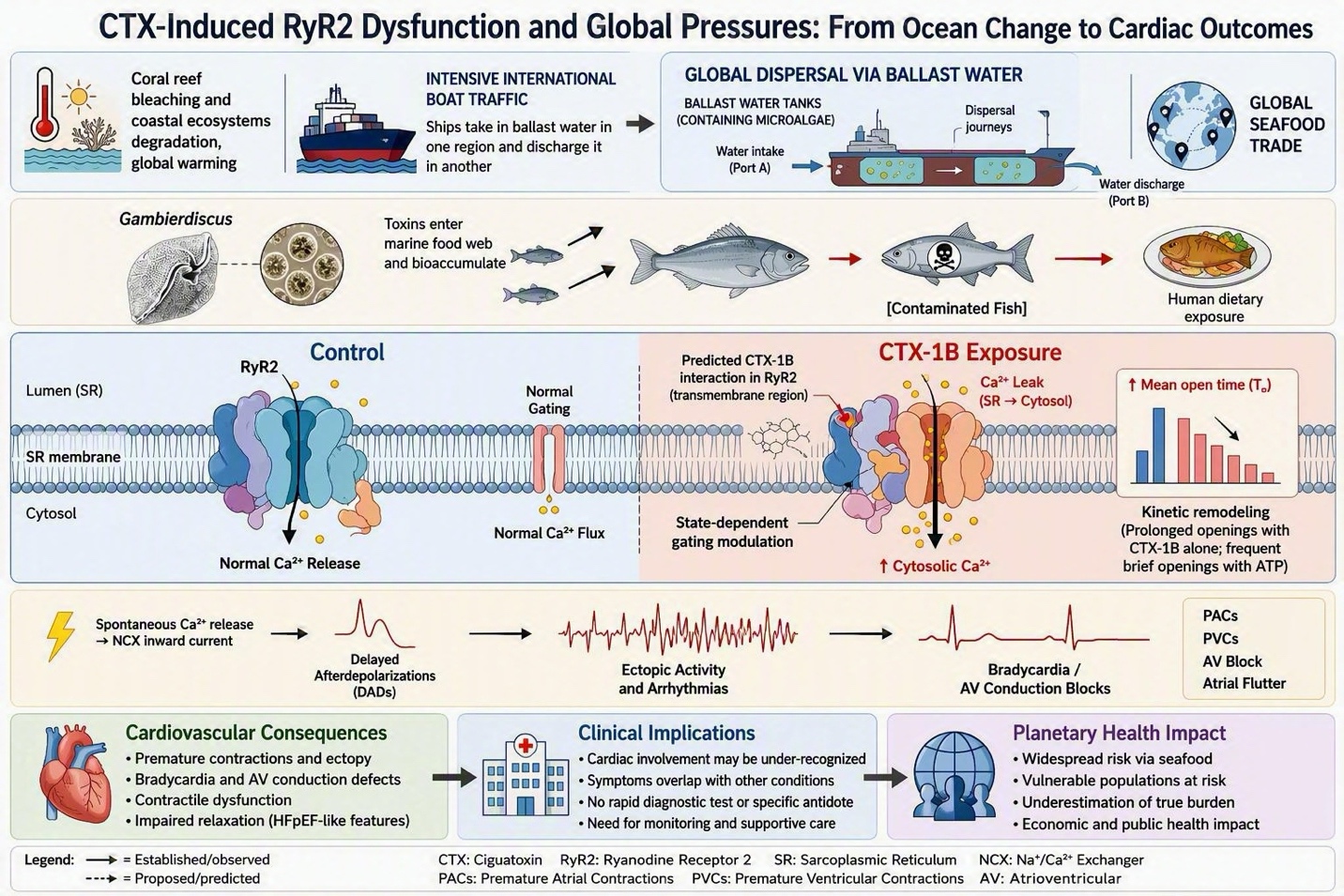
